# Associative Learning and Memory Erasure in a Synthetic Multicellular System

**DOI:** 10.64898/2026.11.09.765282

**Authors:** Jordi Pla-Mauri, Ricard Solé, Núria Conde-Pueyo

## Abstract

Learning is a basic form of adaptation that organisms display from neural systems to single cells. In associative learning, the pairing of two stimuli lets one of them evoke the response to the other. Engineering this behavior in a living cell is difficult, because coincidence detection, memory, and conditional readout must be wired into a single regulatory network. Here we build a consortium of five *E. coli* strains that distributes the operations of associative learning and communicates through one diffusible quorum-sensing signal. The same signal acts as the memory trace. One strain detects the coincidence of the two inputs, a second maintains the memory through positive feedback in the shared medium, a third reads it out on the conditioned stimulus, a fourth reports the unconditioned response, and a fifth erases the memory on command. We show experimentally that the assembled consortium learns the association, since a stimulus that is silent before training evokes a strong response after pairing. Memory is retained through a medium exchange in which both stimuli are removed, and then the conditioned stimulus alone evokes the response. The reporter is not produced without its input. An erase command returns the response to the background. A population model of the consortium accounts for the sequence of responses and gives design rules for writing, retention, and erasure, and a trade-off in storing memory in a shared extracellular signal. Distributing the operations over strains makes each of them separately tunable and testable and provides a route to microbial communities that learn from their history.

## I. INTRODUCTION

The emergence of cognitive complexity is one of the major transitions in the history of life. Alongside innovations in biological individuality and information processing, the evolution of learning provided organisms with the capacity to modify their behavior in response to experience, expanding the range of possible interactions with their environment [1, 2]. Among these capacities, associative learning has been proposed as a particularly important evolutionary innovation, potentially contributing to the Cambrian diversification by enabling new ecological interactions, behavioral flexibility, and coevolutionary arms races [3]. More elaborate forms of associative learning have even been proposed as a key evolutionary transition associated with the emergence of complex cognition [4, 5]. How such capacities first emerged from simpler molecular and cellular mechanisms remains an open question. Synthetic biology offers a complementary strategy to historical reconstruction: by engineering minimal systems that implement specific cognitive functions, we can identify their necessary components and explore how new forms of information processing might arise from simpler biological organizations [6].

Learning is a change in behavior that follows from experience. It defines, along with memory, a crucial precondition for cognition. Its experimental study began with the observation that the consequences of an action alter how often it is repeated, continued with the controlled pairing of stimuli, and led to an analysis of how consequences shape repeated behavior [7–9]. In classical conditioning, a conditioned stimulus (CS) that initially evokes little or no response is presented together with an unconditioned stimulus (US) that already elicits one. After pairing, the CS evokes the response on its own. Later quantitative theories established that the CS must predict the US [10, 11], and which stimuli can be associated depends on the organism [12, 13].

This behavioral definition constrains the physical basis of memory: the response must depend on the relation between the CS and the US. A persistent response to a single stimulus records a history, as in habituation and sensitization, and does not encode a relation between two inputs [14–17]. A minimal associative system therefore needs three functions. It must detect the coincidence of the two inputs, retain a state after they are withdrawn, and allow the CS to retrieve that state when the US is absent. Extinction, the loss of the response after repeated presentations of the CS without the US, leaves the association in place [18]. Here erasure is a separate operation, the removal of the molecular signal that carries the memory. We use an operational definition of learning: a system learns an association if a pairing of two inputs changes its later response to one of them.

In the nervous system, memory has long been attributed to changes in the strength of connections between neurons. Connections were proposed to strengthen when the neurons they join are active together [19], and attractor networks later described recall as convergence to a stable state from a partial or noisy cue [20]. At the molecular level, long-term storage depends on gene expression and on structural change at the synapse [21]. The molecules that hold such a state are continually degraded, which raises the question of how a memory out-lasts its components [22]. Positive feedback provides an answer. A component that promotes its own production can maintain a self-sustaining state that a transient input switches on, as in autophosphorylating kinases and in bistable genetic circuits [23–25]. In cells the feedback must also exceed loss by degradation and by dilution during growth for the high state to persist.

The same questions arise in organisms without neurons. Basal cognition treats learning, memory, and decision making as capacities that predate nervous systems and are implemented by molecular and cellular networks [26–29]. Habituation has been described in the ciliate *Stentor* and in the slime mold *Physarum* [14– 16], experience-dependent changes in responsiveness in plants [30] (see however, [31]), and history-dependent behavior in bacteria [32–34]. In some of these systems the record is kept outside the organism. *Physarum* avoids regions it has already explored by sensing the extracellular slime it left there [35], and termites coordinate nest building through modifications of the environment made by earlier workers, a mechanism known as stigmergy [36]. Microorganisms also anticipate environmental changes through correlations shaped by evolution. In *E. coli*, lactose exposure induces genes used to metabolize maltose, which commonly follows lactose in the gut, and temperature similarly precedes host-associated hypoxia [37, 38].

Regulatory coupling in *Pseudomonas putida* links arsenic exposure to herbicide degradation [39], and laboratory evolution has produced anticipatory responses to sequential cues [40]. Phenotypic switching and signaling that amplifies itself extend cellular responses over time or exploit environmental statistics [41–43]. In all of these cases the association was fixed by selection or adaptation, and none was acquired through a pairing imposed during the experiment.

Synthetic biology has produced molecular records of past events using bistable circuits, DNA inversion, and genomic recording [44–46]. These systems record whether an event occurred. Associative learning additionally requires that the system detect the coincidence of two inputs and then let one of them retrieve the resulting state. An AND gate can detect the pairing [47], feedback can maintain the state, and a second gate can condition retrieval on the CS. Circuits of this kind have been proposed for single cells [48, 49], and a Pavlovian-like conditioning circuit has been built in *E. coli* [50].

Within one cell the rule requires several regulatory interact ions that must be tuned and insulated from crosstalk. Synthetic consortia distribute the operations across strains linked by diffusible signals [51–53]. Quorum-sensing molecules suit this purpose because their shared concentration reports population activity and can communicate a state between cell types [54]. Engineered communities have used diffusible signals for logic, spatial patterning, synchronized oscillation, and collective bistability [55–59]. In this architecture the extracellular signal can also serve as the memory trace, as in the external records above. Any population with the appropriate receptor can read it, an enzyme can degrade it, and a medium exchange removes it. A memory that survives an exchange therefore requires the cells to retain enough signal-producing capacity to restore the trace.

Along with a reliable learning, we also aimed at introducing a predictable meganism for memory erasure. Indefinite memory persistence can be as disadvantageous as the inability to learn; in nature, active memory erasure is a fundamental cognitive and evolutionary requirement to prevent storage saturation, correct obsolete associations in fluctuating environments, and enable behavioral flexibility. Within associative learning, this removal of what was originally learned is called *Pavlovian Extinction* [18]. Within synthetic biology, while previous designs have focused primarily on achieving permanent records via bistable toggle switches or irreversible DNA inversions [25, 45], implementing dynamic reset mechanisms remains a complex and often overlooked challenge. Here, we describe and implement a synthetic circuits capable of the full learning protocol: the acquisition of an association, its retention and, critically, its controlled erasure via a molecular reset operation, demonstrating how distributed cellular cooperation can overcome the constraints of single-cell circuits to achieve reusable memory.

Associative learning has been proposed for two-cell consortia, in which a molecular signal connects the populations and a switch represents the learned state [60]. Those designs were theoretical. Here we build and test a five-population consortium that implements the full protocol: acquisition of an association, its retention after the stimuli are removed and the medium is exchanged, retrieval by the conditioned stimulus alone, and erasure by a separate command. The design assigns coincidence detection, storage, conditioned and unconditioned read-out, and erasure to separate AND, MEMORY, RECALL, DIRECT, and ERASER populations, so that each operation can be tuned and tested on its own.

Flow cytometry of the assembled consortium shows that the conditioned stimulus evokes no response before pairing and a strong response after pairing, that the response is retained through a medium exchange in which both stimuli are absent, and that an erase command returns it to background. A population model of the design gives the conditions for writing, bistability, retention through exchange, and erasure, including finite doses, finite phase durations, and promoter leak. Storing the memory in an extracellular signal imposes a trade-off: the leak bound that keeps the dark state and the gain needed to survive a medium exchange are compatible only when promoter leak is small compared with the square of the fraction of cells retained.

The paper is organized as follows. Section II describes the consortium and the role of each population. Section III develops the population model and derives conditions for writing, retention, and erasure. Section IV presents the measured conditioning and erasure outcomes alongside model predictions. Section V discusses the scope and implications of the results, Sec. VI summarizes the conclusions, and Sec. VII and the Appendices give the experimental methods, model parameters, and promoter calibration.

## II. DESIGN OF THE CONSORTIUM

### A. Five populations, one shared signal

The consortium comprises five *E. coli* populations mixed in a common medium. Each population carries a single plasmid and performs one operation in the conditioning protocol (Table I, Fig. 1). All five share a host background that constitutively supplies the AraC/AraE system, LacI and TetR. Arabinose therefore sets receptor abundance, IPTG gates the *lac*-operator modules, and aTc controls erasure.

**Table 1.** The five consortium populations plus the separate lactone reporter. Consortium strains share a common host back-ground supplying AraC/AraE, LacI^q^ and TetR. Arabinose is fixed within an experiment and sets receptor abundance. The reporter is a TOP10 derivative and is not co-cultured with the consortium.

| Strain | Receptor | Output module | Operation |
|---|---|---|---|
| AND | LasR (pBAD) | *luxI* (pLas-lacO) | write on CS ∧ US |
| MEMORY | LuxR (pBAD) | *luxI* (pLux) | hold by feedback |
| RECALL | LuxR (pBAD) | *gfp* (pLux-lacO) | conditioned response |
| DIRECT | LasR (pBAD) | *gfp* (pLas) | unconditioned response |
| ERASER | — (host TetR) | *aiiA* (pTet) | delete on aTc |
| Reporter (TOP10) | LuxR (J23100) | *gfp* (pLux) | lactone bioassay, not in consortium |

**Figure 1.**
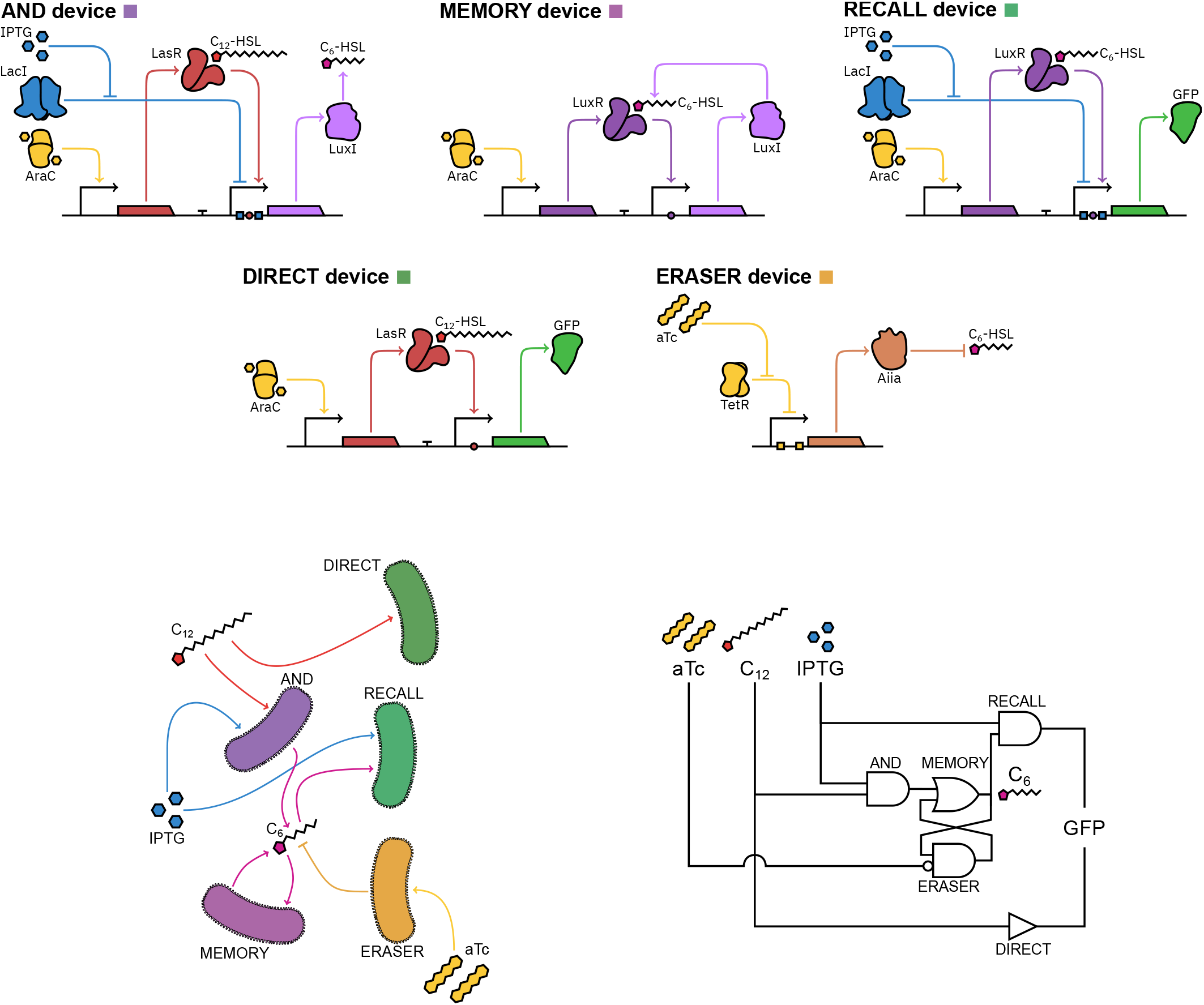
Five synthetic device designs, their shared-medium interactions, and the corresponding circuit. Left, the AND, MEMORY, RECALL, DIRECT, and ERASER strain designs. Center, signaling among the populations through the shared medium: the AND strain writes the 3-oxo-C_6_-HSL signal when IPTG and 3-oxo-C_12_-HSL coincide; MEMORY amplifies and maintains it; RECALL reads it in the presence of IPTG; DIRECT reports 3-oxo-C_12_-HSL; and ERASER produces AiiA in response to aTc to degrade 3-oxo-C_6_-HSL. Lower right, the corresponding circuit-level representation of writing, storage, readout, and erasure.

The AND population performs coincidence detection. Its *luxI* gene sits under a pLas promoter containing a *lac* operator, so LuxI is produced only when LasR is activated by 3-oxo-C_12_-HSL and LacI is relieved by IPTG. LuxI synthesizes 3-oxo-C_6_-HSL, which diffuses out of the cells into the shared medium. The AND population therefore converts the simultaneous presence of the two conditioning inputs into the extracellular memory signal, and neither input alone produces it.

The MEMORY population stores the resulting state. It expresses LuxR and puts *luxI* under pLux control, so 3-oxo-C_6_-HSL drives further 3-oxo-C_6_-HSL synthesis. The feedback loop closes through the extracellular medium and, above a threshold loop gain, supports two stable states. Memory is therefore distributed over two physical pools: the extracellular 3-oxo-C_6_-HSL concentration, which is externally addressable and readable by other populations, and the intracellular LuxI pool retained by the cells. The intracellular pool decays with protein turnover and is the memory that survives immediately after the medium is replaced. Its turnover makes the duration of erasure as important as the eraser dose. The RECALL population implements the conditioned readout. Its *gfp* gene is controlled by a pLux promoter carrying a *lac* operator, so fluorescence appears only when the stored 3-oxo-C_6_-HSL signal and IPTG are both present. Reading is passive: the CS opens the readout gate without modifying the stored state.

The DIRECT population provides the unconditioned readout. Its *gfp* gene sits under a plain pLas promoter, so 3-oxo-C_12_-HSL drives fluorescence directly. This population reports the US response against which the conditioned response is compared.

The ERASER population removes the stored state. It expresses the lactonase AiiA [61, 62] under pTet control, so aTc induces enzymatic degradation of 3-oxo-C_6_-HSL in the shared medium. The enzyme acts on the extracellular signal and does not reach the LuxI retained inside the other populations, so complete erasure requires the degradation of the trace and the loss of the intracellular synthase, which turnover removes and the medium exchanges carry away.

Assigning each operation to a separate strain allows its output to be measured at the module level: the AND strain produces lactone, the MEMORY strain sustains the signal, the RECALL and DIRECT strains report their respective inputs, and the ERASER strain produces AiiA. This division also introduces the strain fractions and the biomass as parameters of the population model. The repressor–operator and arabinose-inducible elements provide independent regulation of transcriptional units in *E. coli* [63]. IPTG serves as the CS because the cells do not consume it during a phase [64].

The classical protocol maps onto the design as follows: the CS is IPTG, the US is 3-oxo-C_12_-HSL, the memory trace is 3-oxo-C_6_-HSL, and the erase command is aTc. Neither input alone activates the AND population, so neither writes memory on its own; their conjunction initiates writing; the CS alone subsequently reads the stored state; and aTc initiates erasure. Crosstalk between the LuxR and LasR systems has been documented in engneered consortia [65]. The doses separate the two channels, and the single-input phases provide the DIRECT and baseline readouts.

#### a. Promoter calibration

The half-response scales fitted to the measured promoter responses define the normalized model inputs; the curves and the fitting procedure are given in Supplementary Material, Sec. S2.

## III. MODEL

### A. Population model

The population model describes signal production, feedback, readout, and enzymatic signal removal. Its reduction yields conditions for writing, retention, and erasure (Table II), which are then extended to finite doses and phase durations, biomass changes, and promoter leak. The model accounts for the sequence of responses of both protocols (Sec. IV) and identifies the parameter combinations that determine each operation.

**Table 2.**
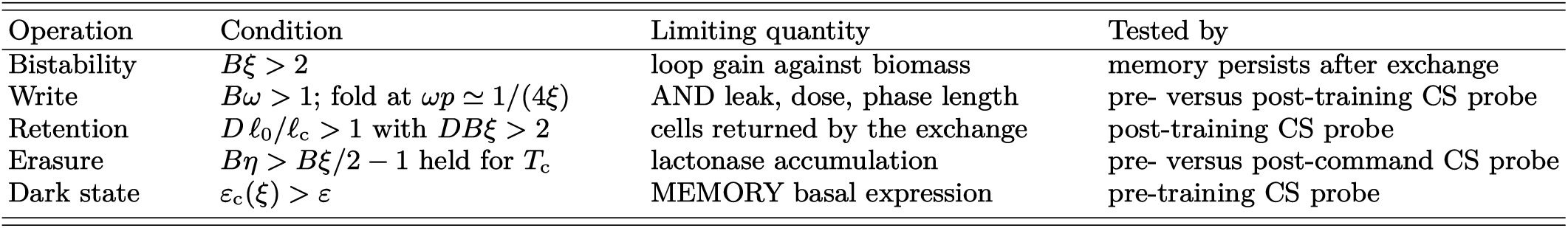
Design rules of the reduced model and the readout associated with each. Every entry is a condition on the dimensionless groups defined in the model; the design point is in Table S1.

| Operation | Condition | Limiting quantity | Tested by |
|---|---|---|---|
| Bistability | Bξ > 2 | loop gain against biomass | memory persists after exchange |
| Write | Bω > 1; fold at ωp ≃ 1/(4ξ) | AND leak, dose, phase length | pre- versus post-training CS probe |
| Retention | D ℓ<sub>0</sub>/ℓ<sub>c</sub> > 1 with DBξ > 2 | cells returned by the exchange | post-training CS probe |
| Erasure | Bη > Bξ/2 − 1 held for T<sub>c</sub> | lactonase accumulation | pre- versus post-command CS probe |
| Dark state | ε<sub>c</sub>(ξ) > ε | MEMORY basal expression | pre-training CS probe |

Because each protein is produced by a single population, the consortium can be described by culture-wide concentrations without resolving cell type. We take the strain fractions to be fixed over a protocol, which amounts to assuming negligible metabolic burden and no growth-rate advantage of any population, and absorb them into effective synthesis rates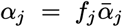. Total biomass *B*(*t*), in OD units, follows the logistic growth of Eq. (1),

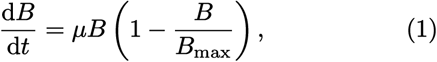

which the reduced model below holds fixed near carrying capacity, *B* ≃ *B*_0_.

The dynamic species are the synthase *L* (LuxI), trace *x* (3-oxo-C_6_-HSL), reporter *G* (GFP), and lactonase *A* (AiiA). The receptors *S* (LasR) and *X* (LuxR) and the repressors LacI and TetR are constitutive and therefore constant at the fixed arabinose concentration used throughout. The inputs are the US *s* (3-oxo-C_12_-HSL), the CS *i* (IPTG), and the erase command *a* (aTc).

We normalize each input by the half-response concentration of its module, *s* = [3-oxo-C_12_-HSL]*/κ*_las_, *i* = [IPTG]*/κ*_lac_, *a* = [aTc]*/κ*_tet_ and *x* = [3-oxo-C_6_-HSL]*/κ*_lux_, with the *κ* given in Supplementary Material, Sec. S1. The reference *x* = 1 is therefore the LuxR half-response, where the conditioned reporter is half open.

With transcription-factor binding fast relative to expression, each promoter is represented by its steady activation with Hill coefficient two,

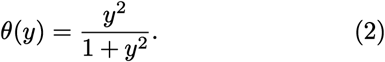

Each gated promoter is approximated by the product of its two single-input responses. Basal expression and crosstalk are set to zero in the protocol model (basal expression is reinstated in Sec. III D), and mixing and signal exchange are taken to be fast enough for 3-oxo-C_6_-HSL to be described by a single shared concentration. The resulting circuit is

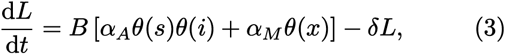

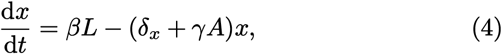

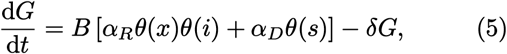

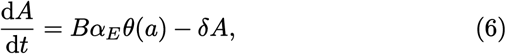

Here *α*_*A*_, *α*_*M*_, *α*_*R*_, *α*_*D*_, and *α*_*E*_ are effective maximal expression rates per OD unit, already weighted by the strain fractions. The parameter *δ* is the protein loss rate, which represents dilution of the protein pools by cell growth and is held constant over a protocol. The parameter *δ*_*x*_ is the passive loss rate of extracellular trace, *β* the trace production rate per unit synthase (it absorbs the culture volume, so that *βL* is a rate), and *γ* the effective trace-clearance rate per unit accessible lactonase.

Equations (3) and (4) describe the memory dynamics, Eq. (5) the one-way readout, and Eq. (6) the erase channel, which enters through the loss term *γAx*.

### B. Reduction to two variables

Two of the four variables decouple. The reporter *G* does not feed back, so Eq. (5) can be evaluated once the memory trajectory is known. The AND and MEMORY populations both produce LuxI, which has a single loss rate, so only the total synthase *L* enters Eq. (3); the two contributions are separated only in the simulated trajectories.

We take the culture to be at constant biomass, *B* ≃ *B*_0_ = 1, so that production does not scale with growth. The pairing drive is kept graded, *p*(*t*) = *θ*(*s*)*θ*(*i*) ∈ [0, 1], so that sub-saturating doses give partial writes; the protocols apply saturating doses, for which *p* is 0 or ≃ 1.

For the eraser we keep the enzyme dynamics, because the duration of erasure matters. Writing the load it imposes as

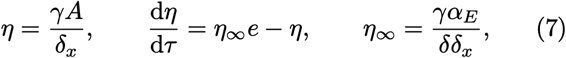

with *e*(*t*) = *θ*(*a*) ∈ [0, 1] the graded erase drive. The saturating aTc dose gives *e* ≃ 1, so the protocols use only *e* = 0 or *e* ≃ 1. A saturating dose held for several turnovers drives *η* to its ceiling *η*_∞_, and a phase long compared with turnover is described by *η* = *η*_∞_*e*. For a shorter phase it is the accumulated enzyme, not the dose, that sets how long erasure lasts.

Measuring the synthase as the trace it sustains and time in protein turnovers,

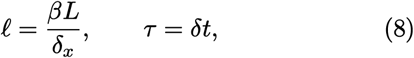

turns the two remaining equations (with d*/*d*τ* = *δ*^−1^ d*/*d*t*) into a two-dimensional system for the memory, Eqs. (9) and (10) below, in which *θ* is the common response function of Eq. (2),

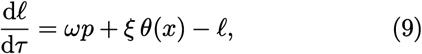

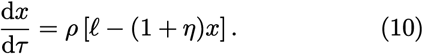

Here *ℓ* is the trace the synthase would sustain with the eraser off: at steady state *ℓ* = (1 + *η*)*x*, and with *η* = 0 the two coincide.

The reduced model is therefore described by four dimensionless groups.

The *write capacity*

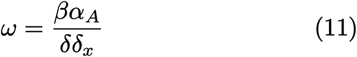

is the trace the AND module sustains at saturation, expressed in the half-response units of *x*.

The *memory gain*

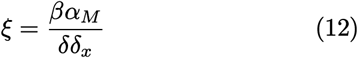

is the gain of the MEMORY loop. With the write off it alone determines whether that loop is bistable: one fixed point exists for *ξ <* 2, and three exist for *ξ >* 2.

The *saturated erase load η*_∞_ of Eq. (7) measures lactonase clearance relative to passive trace loss, with *η*(*t*) = *η*_∞_*e*(*t*) the load once the enzyme has reached steady state. It divides the write and the loop gain by the same factor, giving the effective loop gain

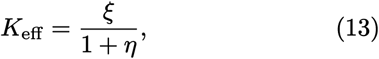

which reduces to *K*_eff_ = *ξ* with the eraser off. The load is the only one of these groups that the protocol varies.

Finally, the *recovery-speed ratio*

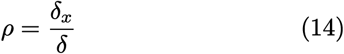

rescales the trace equation. It enters no fixed point, so it does not affect which states exist or their values, but it does enter the retention threshold *ℓ*_c_ of Eq. (18), which is a location in state space.

Away from *B* = 1 the effective gains scale with biomass (*Bω, Bξ, Bη*_∞_), so every rule below is a condition on these products, and bistability requires *Bξ >* 2. A culture diluted below *B* = 2*/ξ* therefore has no ON state to hold, and recovers one only as it grows back toward saturation. The write is gated in the same way: the fold of the paired drive is crossed only above 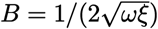, which lies below 2*/ξ* whenever *ξ <* 16 *ω*. In the window 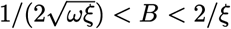 the write can switch the loop although the loop is not bistable, and below the lower density neither the write nor the feedback has any effect. The protocol resuspends the culture at *B*_stim_ (Table S1), below both densities. Every active phase therefore opens with the circuit off, and nothing is written or held until growth carries the culture back across the fold. The conditions below are evaluated at the density of interest, and the scans are computed at *B* = 1.

### C. Write, retention and erasure

Setting the derivatives of Eqs. (9)–(10) to zero gives

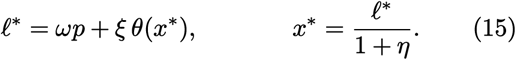

The eraser divides the write and the feedback gain by the same factor. With the write off, *p* = 0, the steady states therefore depend on *ξ* and *η* only through *K*_eff_ of Eq. (13). In that case *x*^*^ = 0 is always stable, and a pair of additional solutions appears at the fold *K*_eff_ = The value two is geometric: nonzero solutions satisfy *K*_eff_ *θ*(*x*)*/x* = 1, and *θ*(*x*)*/x* = *x/*(1 + *x*^2^) peaks at *x* = 1 with height one half, so the feedback crosses the line of unit slope only when its gain exceeds the reciprocal of that peak. The pair appears at *x* = 1, the LuxR half-response. The larger solution is stable and the smaller a saddle, and for large gain the ON branch approaches *K*_eff_ from below. In closed form the fixed points are 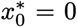 and, for *K*_eff_ ≥ 2,

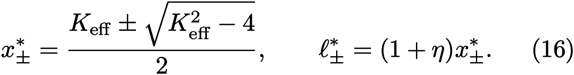

Bistability alone does not guarantee that the memory is set. A resting culture begins at *x* = 0, and switching requires a perturbation large enough to carry the state past the smaller of the two solutions of Eq. (16). Here that perturbation is the simultaneous application of the US and CS, *p* = 1.

The three circuit operations are *write*, which establishes the state; *retention*, which lets it survive removal of the inputs; and *erasure*, which removes the ON state on command. Each gives a design rule on the four groups.

Finite protocols must satisfy these rules at a given dose, duration and biomass, which Sec. III D takes up.

#### a. Write

With the inputs off the loop rests at *x* = 0. The paired inputs add the drive *ωp*, which lifts that rest point until, at a fold *w*_*_, it meets the saddle and both disappear, leaving the ON state as the only solution. With the eraser off, the fold is the double root of Eqs. (15), at *w*_*_ ≃ 1*/*(4*ξ*), so a pairing *p* writes when 4*ξ ωp >* 1. We adopt the stricter rule

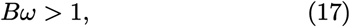

which ensures that the write alone carries the trace to the readout reference *x* = 1 with the loop open, so that writing does not presuppose a closed loop. The condition is on the write capacity and is independent of the bistability condition *Bξ >* 2. Inside the window 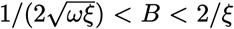 the write can switch the loop but cannot lift the trace to *x* = 1, since *Bω <* 2*ω/ξ* ≤ 1 there for *ξ* ≥ 2*ω*. Equation (17) therefore applies to a circuit that is already bistable, and the fold alone governs the window. Below 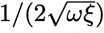 the fold is out of reach at any dose and no state can be established.

#### b. Retention

Once the memory has been written, the inputs can be removed while the cells are retained, by centrifuging and washing with new medium. A wash empties the trace from the medium but returns a fraction of the cells, and of the proteins they carry, to fresh medium. The culture restarts at *x* = 0 carrying only the synthase left by the pairing, *ℓ*_0_ = (1+*η*)*x*^*^ = *ωp*+*ξ θ*(*x*^*^). With the load withdrawn, integrating Eqs. (9)–(10) from this state shows that the memory is recaptured when *ℓ*_0_ exceeds a critical value *ℓ*_c_(*Bξ, ρ*), the height at which the stable manifold of the saddle meets *x* = 0. Retention therefore imposes two independent conditions. The wash must leave the culture bistable, *DBξ >* 2, or no such threshold exists and no quantity of retained synthase can hold the state. Within the bistable region the retained synthase must exceed the threshold,

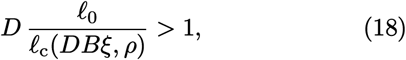

where *D* is the fraction of the biomass and of the intracellular pools that the exchange retains (*D*_wash_ in Table S1), and *ℓ*_c_ is evaluated at the post-wash density *DB*, the density at which the trace has to be rebuilt. The subsequent reset to *B*_stim_ takes the circuit below the fold again, so the state crossing it is carried rather than rebuilt, and Eq. (18) is not evaluated there.

Retention is the requirement specific to storing the memory outside the cells: the wash keeps the synthase and discards the trace, and the loop rebuilds the trace from the synthase alone. An exchange delivers *D ℓ*_0_, with *D* set by the medium handling. The post-exchange CS probe tests this condition directly, because a bright response requires the trace to have been restored.

#### c. Erasure

While the load is present, the largest trace the loop can sustain is *K*_eff_ itself, so at biomass *B* the ON state disappears once

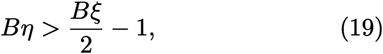

with *η* the load accumulated so far. The load rises only as fast as the enzyme accumulates, so a saturating dose does not erase at once: the ON state persists while the load is below the threshold. Erasure therefore requires a sufficient duration as well as a sufficient dose. The load must first cross Eq. (19) and then remain above it until turnover has carried the synthase below *ℓ*_c_(*Bξ, ρ*). If aTc is withdrawn before that, the restored loop recaptures the ON state from the remaining synthase.

The erase phase is thus a trajectory through the bifurcation diagram (Fig. 2b). The lactonase accumulates according to Eq. (7), and as *η* increases *K*_eff_ falls through the fold and the trace collapses. When the erasure input is withdrawn the gain returns toward its original value, but the trace does not recover, because the remaining synthase cannot carry the system across the saddle of the restored loop.

**Figure 2.**
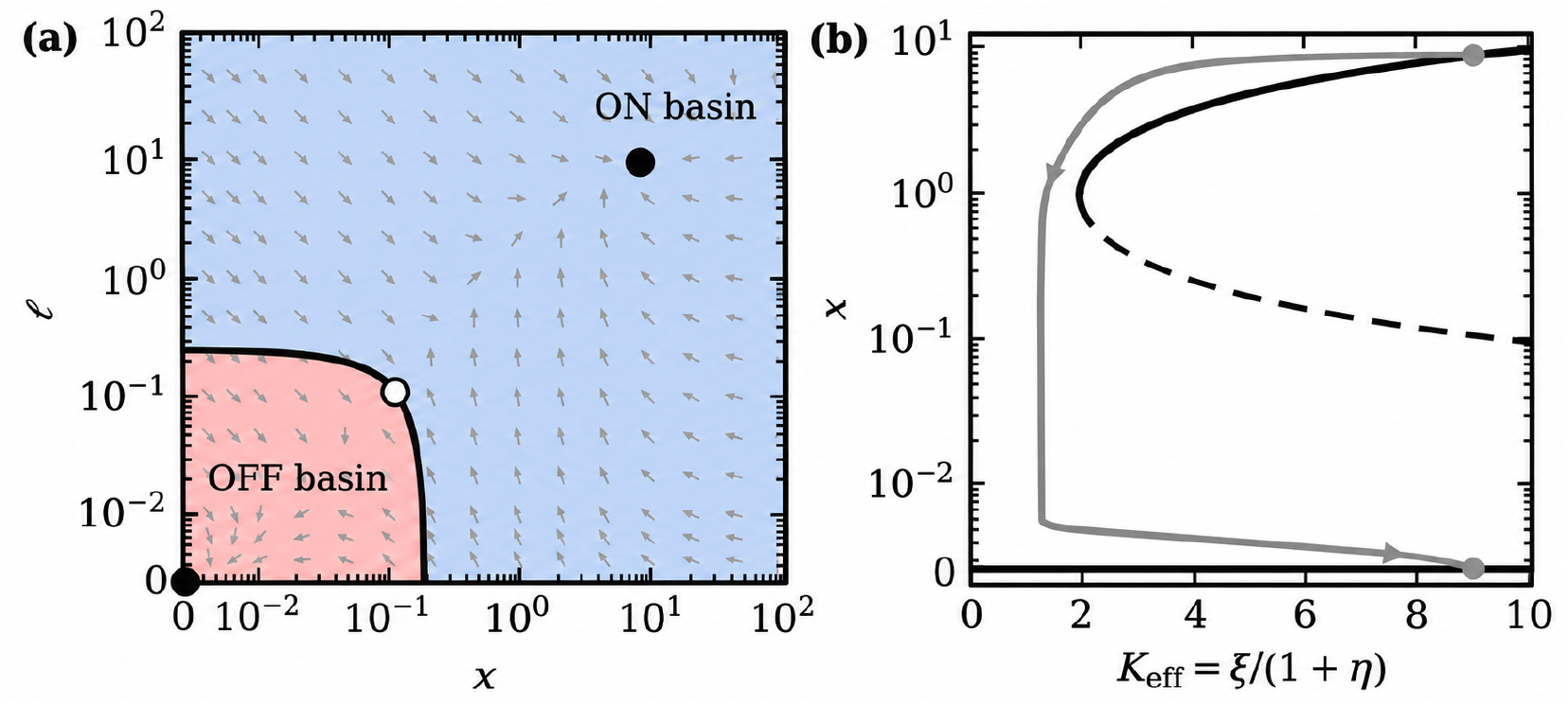
Mechanism of the reduced model, Eqs. (9)–(10). (a) Phase plane on symmetric-logarithmic axes showing both attractors, the ON (blue) and OFF (red) basins, and the vector field (gray arrows). The boundary between the basins is the stable manifold of the saddle (open circle) and meets *x* = 0 at *ℓ*_c_, so a wash is survived when *D ℓ*_0_*/ℓ*_c_ *>* 1 (Eq. (18)). The panel is drawn at *B* = 1; the retention condition evaluates *ℓ*_c_ at the post-wash gain *DBξ*, where the threshold is higher. (b) Bifurcation diagram against *K*_eff_ (black) carrying the gray erase trajectory: load accumulates, gain falls past the fold, trace collapses, and withdrawing aTc returns the gain without returning the state.

### D. Design margins: dose, duration and leak

The three rules above are conditions for the groups: the writing in *Bω*, the retention in *D ℓ*_0_*/ℓ*_c_ together with *DBξ >* 2, and the erasure in *Bη* held past the fold. An experiment fixes a concentration and a phase length and uses promoters that are never fully silent, so each rule must be restated as a dose floor, below which no phase length works, and, for erasure, a minimum duration.

For the write, the floor is the fold itself *Bω p >* 1*/*(4*Bξ*); above it, raising the dose shortens the needed phase. For erase, the floor is the dose condition of Eq. (19) at a saturating command, *θ*(*a*) *>* (*ξ/*2 − 1)*/η*_∞_, which requires *η*_∞_ *> ξ/*2 − 1. Above this floor, erasure is limited by time alone: the load must be held past the fold for a critical duration

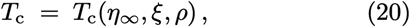

so that turnover carries the synthase below *ℓ*_c_(*Bξ, ρ*). *T*_c_ is measured in units of protein turnover time *δ*^−1^ and diverges as the load approaches the fold above. These margins are computed at *B* = 1, whereas the protocol sweeps the density. The erase duration and the retention ratio are the two margins that set the operating window of the protocols.

#### Leak on the write side

Leak from the AND module is the false-positive risk that the pre-training CS probe is designed to exclude. With IPTG saturating and 3-oxo-C_12_-HSL absent, the fitted pLas basal fraction *ℓ*_las_ (Fig. S1a) contributes a drive *ω θ*(*i*) *ℓ*_las_, which must stay below the fold 1*/*(4*ξ*), that is

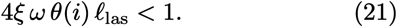

The leaks of the RECALL and DIRECT readouts similarly set the OFF floors of the two CS probes. The trajectories of Sec. IV are computed with basal expression at zero, so none of these floors appears in them.

#### Basal expression

A MEMORY promoter that fires at a fraction *ε* of its ceiling with no trace bound adds a drive to the loop that does not switch off. Its fixed points then solve

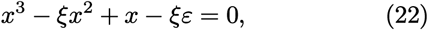

so the dark state is no longer *x* = 0 but a small positive root that rises with *ε* until, at a fold, it meets the saddle and both vanish. Beyond that leak only the ON state remains and the memory cannot be dark. The double root of Eq. (22) and the leak that produces it are

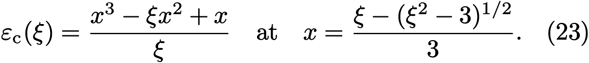

The fold is real for 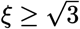, where the discriminant of the derivative of the left-hand side of Eq. (22), 3*x*^2^ − 2*ξx* + 1, changes sign. For *ξ* ≫ 1 the tolerated leak falls as *ε*_c_ ≃1*/*(4*ξ*^2^). A dark state therefore requires *ε < ε*_c_(*ξ*), which is an upper bound on the gain, and lowering the gain to satisfy it also lowers the retention margin.

The dark-state and retention requirements conflict. Because *ε*_c_ decreases with *ξ*, the dark state requires *ξ* below the root of *ε*_c_(*ξ*) = *ε*, whereas bistability after the wash requires *ξ >* 2*/*(*DB*_max_). The two are compatible only if *ε < ε*_c_(2*/DB*_max_), which for *DB*_max_ ≪ 1 reduces to

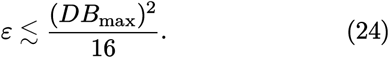

Equation (24) is the compatibility condition for the two bounds: when it holds, a range of gains can preserve both the dark state and the post-exchange ON state. It expresses the trade-off of extracellular storage: promoter leak must be low relative to the square of the retained cell fraction. Lower leak or greater cell retention widens the compatible gain range.

The model treats extracellular signal removal and cell retention as independent. If retaining more cells also retains more signal, Eq. (24) depends on both fractions.

## IV. RESULTS

### A. Learning, retention, and erasure in the assembled consortium

The consortium learns the association, retains it through a medium exchange, and loses it on command (Figs. 3 and 4). In the conditioning protocol the US-alone phase gives the unconditioned response through the DIRECT population. The CS probe that follows is at background, so neither the US phase nor the CS input has written a memory. Paired exposure is the brightest phase. The CS probe after pairing, delivered after the US has been withdrawn and the medium exchanged, is bright and comparable to the US-alone phase. The same input that produced no response before pairing produces the response after it. The comparison across protocols has the same outcome at matched positions in the schedule: the second phase is a CS probe in both, dark in the conditioning protocol and bright in the erasure protocol, where pairing precedes it. The pattern is the same in all three independent cultures.

**Figure 3.**
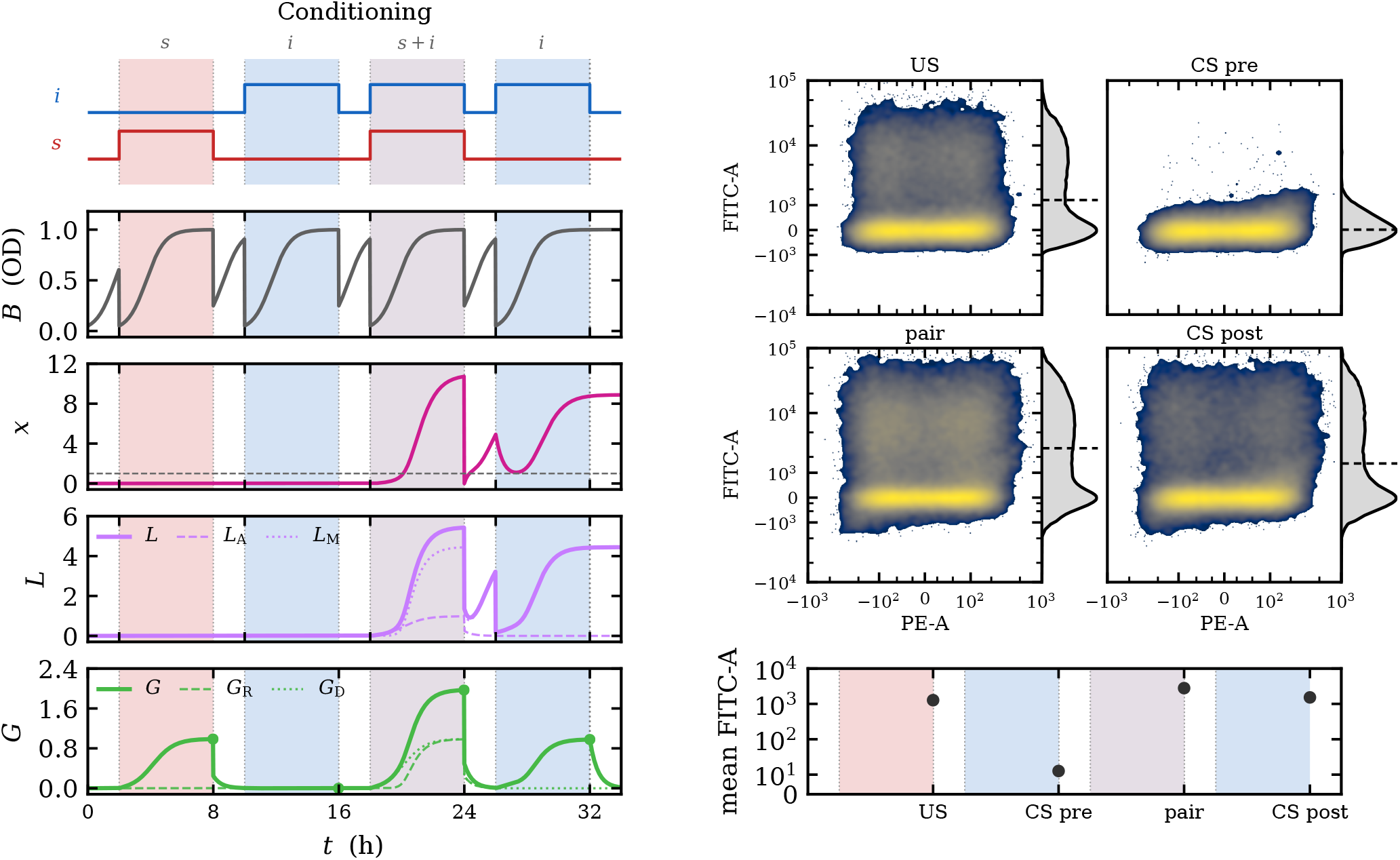
Conditioning. Left, modeled protocol trajectories: inputs, biomass *B*, trace *x*, synthase *L* with its AND and MEMORY contributions, and reporter *G* with its RECALL and DIRECT contributions, with the phase-end sampling points marked. Right, measured single-cell fluorescence at the end of each active phase, as FITC-A against PE-A density maps with FITC-A marginals, pooling the three independent cultures. The lower panel shows the pooled mean FITC-A for each phase; the three culture means differed by less than 5% from one another within each condition. The pre-training CS probe is at background while the post-training CS probe, delivered after the US has been withdrawn, is bright; that pair is the conditioned response.

**Figure 4.**
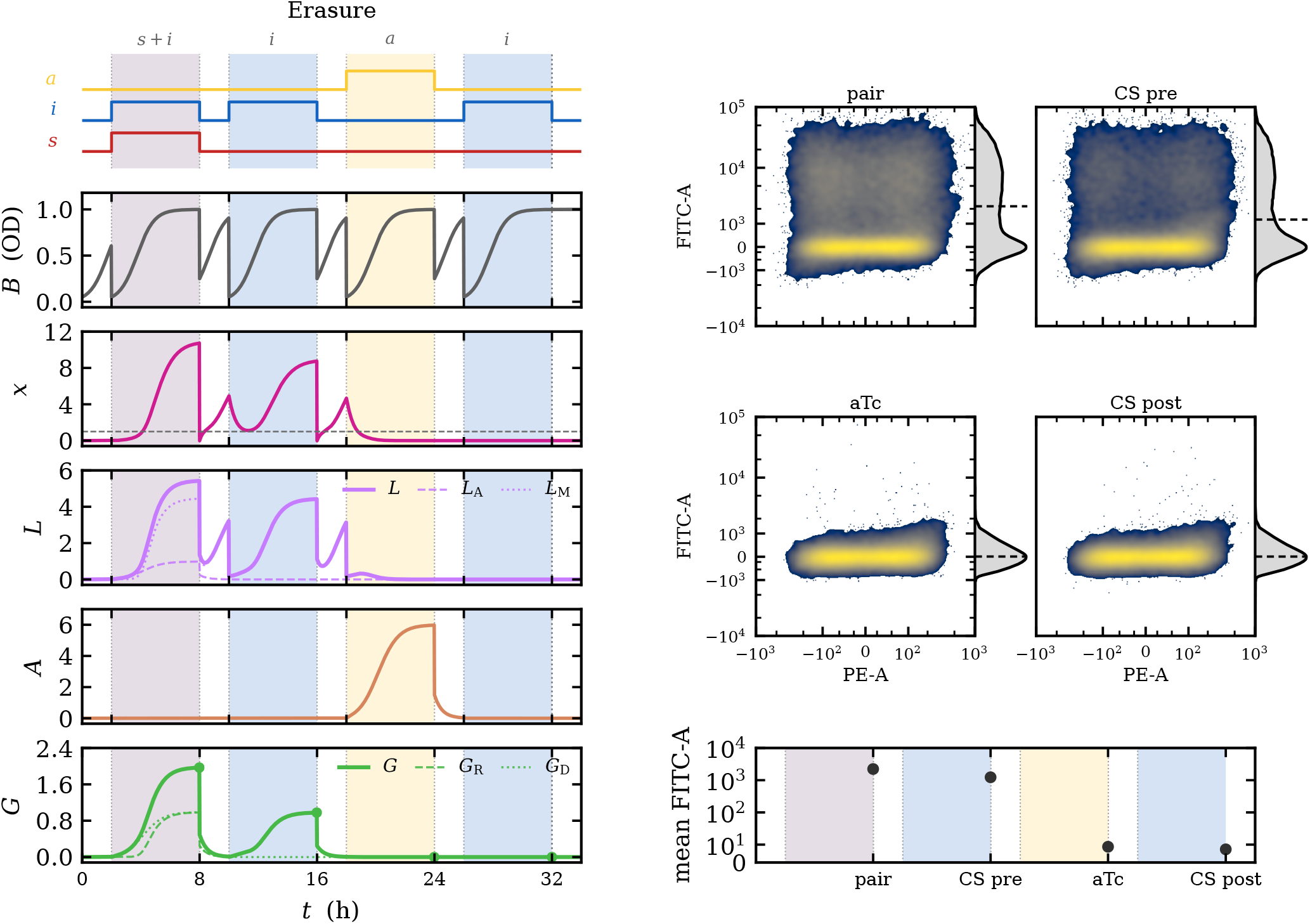
Erasure. Left, modeled protocol trajectories, with the lactonase pool *A* added: the aTc phase drives *A* up, the trace *x* falls below the recovery threshold, and the synthase pool does not rebuild it. Right, measured single-cell fluorescence at the end of each active phase, plotted as in Fig. 3. The lower panel shows the pooled mean FITC-A for each phase; the three culture means differed by less than 5% from one another within each condition. The CS probe before the command is bright and the probe after it returns to background, at the level of the pre-training probe of the conditioning protocol.

The erasure protocol gives the complementary result.

Its first two phases reproduce the write: pairing is bright and the CS probe that follows remains bright. After the aTc phase the CS probe returns to the dark floor, at the level of the pre-training probe of the conditioning protocol. The erase command removes a memory that the preceding probe shows to be retained.

The reporter does not build up between phases. The sample taken after the aTc phase, which follows a bright probe and a wash, is at background, and the CS probe after the US-alone phase is dark, so GFP from one phase does not carry over and the reporter is produced only in phases that supply its inputs. The medium exchange clears the extracellular signal, and IPTG is the only input applied in the post-training probe. The response to IPTG alone therefore requires signal made after the exchange by the retained cells. The same probe is dark when no pairing precedes it and after the erase command.

The bright phases are bimodal: a bright subpopulation coexists with cells at the background level, and the mean FITC-A reflects the bright tail. The dark phases lie at the autofluorescence floor.

### B. Model account of the protocols

The mean-field model gives reporter production at each phase from the gate states through

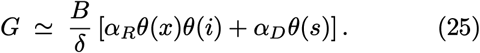

The conditioning schedule visits all four combinations of the two gates. US alone opens DIRECT; CS alone before training opens neither gate; pairing opens both; and the post-training CS probe opens RECALL alone. The erasure schedule gives a bright pre-command CS probe followed by a dark post-command probe. These are the sequences measured in Figs. 3 and 4, whose left panels show the corresponding model trajectories.

The model also accounts for what happens between the sampled endpoints. During a wash no input is present, so RECALL and DIRECT are closed and no reporter is produced; the GFP carried over from the previous phase decays with protein turnover and does not contribute to the next probe. The extracellular trace behaves differently. A medium exchange clears it, and when a memory has been written the retained synthase rebuilds it during the wash and the feedback loop returns it to the ON state. When no memory has been written, or erasure has removed the synthase, nothing is rebuilt. The bright post-training CS probe therefore reports a trace that was restored after the exchange, and the dark post-command probe reports one that was not. A single design point of the model gives the observed ordering of the phases in both protocols: pairing is brightest, and the US-alone and post-training CS phases are comparable. Resuspension below the bistable density causes a transient loss of trace, after which growth restores the loop gain and the state is recovered from the retained synthase.

## V. DISCUSSION

Learning and forgetting are fundamental components of cognition, allowing biological systems to acquire, retain, retrieve, and discard information about their environment [21]. Here, we have shown that a synthetic multicellular system can acquire an association between two stimuli, retain and retrieve the resulting memory, and erase it on command. Beyond the implementation of these functions, our results illustrate how elementary cognitive operations can be distributed across specialized cell populations and integrated through intercellular communication. This provides a synthetic approach to investigating the emergence of multicellular cognition: by constructing and manipulating the mechanisms of learning and memory erasure, we can explore the minimal organizational principles that make these capacities possible and how they might be extended toward more complex forms of collective information processing.

The consortium implements each element of associative learning in a separate strain and carries each protocol through in a single culture. A stimulus that is silent before training evokes the response after pairing, the response survives a medium exchange in which both stimuli are withdrawn, and the erase command removes it. The reporter stays dark before pairing and does not carry over between phases. The medium exchange removes the extracellular signal, so the response to IPTG alone after it shows that the retained cells rebuild the trace when a memory has been written, and the stored state is held by the community.

The model places the stored state in two pools: extracellular 3-oxo-C_6_-HSL and the intracellular LuxI that produces it. A medium exchange removes the extracellular pool, and the retained synthase restores it. AiiA-mediated erasure succeeds when signal clearance and synthase turnover together carry the system below the recovery threshold. Persistence and erasure therefore depend on biomass, cell retention, protein turnover, and enzyme accumulation.

The main design constraint is the trade-off between a dark state and retention. Basal expression of the MEM-ORY promoter removes the low state above a leak *ε*_c_(*ξ*) that falls with the gain, whereas bistability after the wash requires a gain of at least 2*/*(*DB*_max_). Both requirements can be satisfied only when Eq. (24) holds. This condition states how small promoter leak must be relative to the retained cell fraction for an operating range to exist.

Associative learning in a two-cell consortium has been analyzed theoretically [60]. The present work realizes the learning protocol experimentally in a five-population consortium, and its model extends that analysis to finite stimulation, biomass changes, promoter leak, and signal removal. Because each operation resides in its own strain, a module can be retuned or replaced without redesigning the others, and the same architecture can be extended with further inputs or readouts. Storing the trace in a diffusible signal makes it addressable by any population with the appropriate receptor, and erasable by an enzyme, which is the property that distinguishes this memory from those held inside a single cell.

Several measurements would extend these results. The fraction of cells returned by the medium exchange and the signal carried over in the resuspended volume would fix the retention condition, and a standard curve for the lactone reporter in matched medium would place the trace in concentration units. An aTc-free run of the erasure schedule would give a matched control for the elapsed time, and a longer schedule would show how long erasure lasts after residual AiiA activity ends. Measuring the basal expression of the MEMORY promoter in the assembled consortium, together with dose–response measurements in the paired state, would test the leak bound and the saturating-dose assumption directly.

More broadly, our results suggest a path toward engineering multicellular systems with increasingly sophisticated forms of information processing. The modular organization of the consortium makes it possible to extend the present architecture with additional inputs, feedback loops, and interacting memories, potentially enabling more complex forms of collective learning and adaptive behavior. Such designs could eventually be incorporated into spatially organized synthetic tissues and organoids [66], or into engineered living constructs such as Xenobots and Anthrobots [67, 68], where sensing, memory, and behavior could become coupled to morphogenesis and coordinated cellular activity. These possibilities remain to be demonstrated, but they point toward a broader experimental strategy for exploring the origins of multicellular cognition. In the spirit of synthetic evolutionary transitions [6], constructing minimal systems that learn, remember, and forget provides a way to investigate how elementary cellular interactions can give rise to new levels of collective organization and cognitive agency.

## IV. CONCLUSIONS

We have built and tested a synthetic *E. coli* consortium in which associative memory is distributed over five populations. The consortium acquires an association on paired input, retains it through a medium exchange, retrieves it on the conditioned stimulus alone, and loses it on an erase command. The model places the stored state in a shared extracellular signal and the enzyme that sustains it. Writing, retention, and erasure reduce to conditions on the write capacity, memory gain, saturated erase load, and biomass; finite doses and phase durations enter as margins on those conditions. The dark state and retention through a medium exchange bound the memory gain from opposite sides, and both hold when the promoter leak is small compared with the square of the retained cell fraction.

## VII. METHODS

### A. Strains and plasmids

*E. coli* was used because its repressor–operator and arabinose-inducible elements allow several transcriptional units to be regulated independently in one host [63], and because the LuxR/LuxI and LasR lactone systems are the best-characterized quorum-sensing channels for communication between engineered populations [53, 54, 65]. The consortium comprised five engineered *E. coli* populations, each carrying a single plasmid (Table I). The common host background was an XL1-Blue derivative bearing an HK022-integrated *araC* – *araE* –*lacI* ^q^ cassette and *tetR* on the F^′^ element [69, 70], which supplies arabinose-inducible expression of the pBAD-driven receptors, constitutive LacI to gate the IPTG-responsive modules and constitutive TetR to control the aTc-inducible eraser. Placing the regulators in the host and the modules on plasmids means that the five strains share every regulator and differ only by one plasmid, and hence by one operation, so a difference in behavior can be traced to a module. A separate TOP10-derived lactone reporter carrying a high-copy-number J23100– *luxR*, pLux–*gfp* construct assayed 3-oxo-C_6_-HSL bioactivity in the medium recovered from centrifuged AND and MEMORY cultures; it was not part of the consortium (Table I). It exists because the memory signal is a small molecule that no cytometer sees directly. DNA fragments were custom synthesized (Twist, GeneWiz), cloned using restriction enzymes and ligation or IVA cloning [71], and verified by sequencing (Plasmidsaurus).

### B. Media, growth and consortium assembly

Glycerol stocks were streaked onto LB agar with chloramphenicol and grown overnight at 37 °C. Single colonies were inoculated into selective medium and grown overnight, diluted into fresh medium, and the five consortium strains combined at equal ratio, unless other-wise specified, to form the culture. All growth, in the pre-cultures and in every phase of both protocols, was in LB with 1% glucose, chloramphenicol and 0.1 mmol L^−1^ arabinose, at 37 °C with shaking at 250 rpm.

LB is not a defined medium, and its composition changes as a culture consumes its carbon and amino-acid sources [72]. Every wash therefore begins from fresh medium after an exchange, and every active phase from the medium of the preceding wash, so that phases are compared under the same starting conditions. Glucose strengthens LacI repression of the *lac*-operator modules, which lowers their basal expression and protects the dark state. Arabinose sets the abundance of the pBAD-driven LasR and LuxR receptors and is held at one low concentration throughout, in the pre-cultures and in every phase; the weakly induced pBAD provides the modest receptor abundance required. The arabinose level therefore enters the effective maximal rates 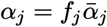 of Sec. III as a constant and not as an input, whereas the half-response scales *κ* are properties of the promoters and do not depend on it. Selection is maintained in every phase. The strain fractions *f*_*j*_ and the biomass are not measured during a protocol and are treated as model parameters.

The three inputs are 3-oxo-C_12_-HSL (the US), IPTG (the CS) and aTc (the erase command), applied at the doses of Table S1. Each dose saturates its own module, *θ* ≥ 0.99 at the fitted half-response scale, so that a phase is defined by which inputs are present and the normalized drives *p* and *e* are approximately zero or one. The doses were chosen from the module curves of Supplementary Material, Sec. S2.

### C. Conditioning and erasure protocols

The protocols translate a behavioral experiment into culture conditions. In conditioning, a baseline probe with the CS alone establishes that the CS does not yet elicit the response, a US-alone phase shows the unconditioned response, pairing supplies the association, and a second CS probe tests it. In erasure, a written state is probed, removed by command, and probed again. Both protocols use the same CS input before and after the manipulation, so that the comparison is made within one culture history.

Each protocol consists of a two-hour wash followed by four six-hour active phases separated by two-hour washes, for a 32-hour schedule (Table III). Inputs are present throughout each active phase and absent from every wash interval. A six-hour phase spans several protein-turnover times, and the two-hour wash that follows each exchange allows the retained cells to recover extracellular 3-oxo-C_6_-HSL. The four phases cover the input combinations of the conditioning and erasure schedules.

**Table 3.**
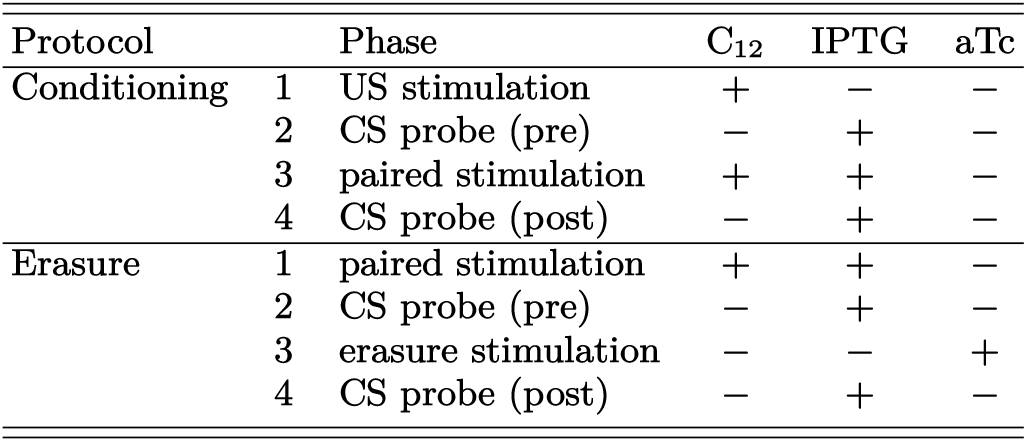
Active-phase sequences of the two protocols. Each protocol has four six-hour active phases, each preceded by a two-hour no-input interval (32 h total). Each later interval began with a medium exchange. In the model, this clears inputs and extracellular trace while retaining a fraction of the cells and intracellular proteins. + denotes the applied dose and − denotes no input.

Each interval between active phases begins with a medium exchange: the culture is centrifuged twice, washed, and resuspended in fresh medium carrying arabinose and selection but none of the three inputs, and the resuspended culture is diluted fourfold. This step is the experimental counterpart of the exchange in the model, which clears extracellular 3-oxo-C_6_-HSL while retaining a fraction *D* of the biomass and of the intracellular LuxI, GFP and AiiA, set by the fourfold dilution to *D*_wash_ = 1*/*4 (Table S1), and which enters Eq. (18) through that fraction alone. It is the step that distinguishes a memory held by the cells from one held by the medium. The cell loss in the centrifugation steps and the 3-oxo-C_6_-HSL carried over in the resuspended volume were not measured and enter the model as parameters. The model treats them as independent, which corresponds to concentrating the cells and discarding the supernatant. A deeper exchange removes more trace but returns fewer cells, so the retention condition is expressed in terms of *D*.

The diluted culture is incubated for the two-hour wash and is then reset to OD 0.05 (*B*_stim_ in Table S1). For the reset the culture is centrifuged and diluted in LB, the volume needed is taken and centrifuged, and the cells are resuspended in the medium in which they grew during the wash, which retains residual 3-oxo-C_6_-HSL, with the inputs of the next phase. The reset sets the cell density without removing the extracellular 3-oxo-C_6_-HSL, as in the model.

*Conditioning* comprises 3-oxo-C_12_-HSL alone (the US), IPTG alone (a pre-training CS probe), 3-oxo-C_12_-HSL with IPTG (paired training), and IPTG alone (a post-training CS probe). *Erasure* comprises paired training, a CS probe, aTc alone, and a final CS probe. The schedules place the same CS input before and after pairing or erasure, with the US-alone phase specifying the DIRECT readout condition.

### D. Sampling and flow cytometry

Flow cytometry resolves cell-to-cell variation that a bulk fluorescence measurement would average, and shows whether bright cells coexist with a dark subpopulation.

Three independent cultures were run through each protocol. Samples were collected from each mixed consortium at the end of each active phase, before the next medium exchange, and analyzed by flow cytometry. Events were recorded in compensated FITC-A and PE-A channels. Compensated values are retained as recorded, including negative values near background, and are displayed with a signed inverse-hyperbolic-sine transform. Summaries are computed from the untransformed values. The density maps pool the events of the three cultures, and the summary panels show the mean of each culture. The event distributions characterize single-cell responses within each sample; individual cells are not additional biological replicates. The three independently run cultures per protocol are the biological replicates.

### E. Lactone bioassay

The TOP10 reporter of Table I grows in the medium recovered by centrifuging AND and MEMORY cultures, and its fluorescence reads the lactone those modules produced. This gives the responses of the lactone-producing promoters in Supplementary Material, Sec. S2. The reporter is a separate culture and not a member of the consortium. Lactone readings are relative to the sensor, and the normalized trace *x* of Sec. III is defined against the LuxR half-response.

## ACKNOWLEDGMENTS

The authors thank Paul Cozmuta and the members of the Complex Systems Lab for useful discussions. This work was supported by AGAUR FI-SDUR 2020 and grant AEI-PID2023-152129NB-I00. JPM was supported by grant PRE2020-091968, funded by MCIN/AEI (10.13039/501100011033) and co-funded by the European Social Fund through the program *Investing in your future*. RS and JPM also thank the Santa Fe Institute, where part of this work was developed.

## AUTHOR CONTRIBUTIONS

J.P.-M. designed and constructed the strains and plasmids, performed the experiments and the cytometry, carried out the model analysis and simulations, and wrote the manuscript. N.C.-P. conceived the original idea and the base design, designed plasmids, and performed cytometry. R.S. supervised the study, acquired funding, and wrote the manuscript. All authors contributed to the conceptualization, discussed the results and revised the manuscript.

