## Supplementary Material for "Associative Learning and Memory Erasure in a Synthetic Multicellular System"

#### S1. MODEL PARAMETERS

The inputs to the model have three sources (Table S1). The half-response scales $\kappa$ are fitted to the measured promoter responses of Appendix S2. The doses and phase durations are those of the protocols. The remaining entries define the design point at which the model trajectories of Figs. 3 and 4 and Fig. 2 are drawn. The rules of Sec. III depend on the rates only through the dimensionless groups $\omega$, $\xi$, $\eta_\infty$ and $\rho$. Each protein is produced by a single population, so the effective synthesis rates are $\alpha_j = f_j\bar{\alpha}_j$ for a strain fraction $f_j$ and a per-biomass rate $\bar{\alpha}_j$, and a change in consortium composition changes the effective rate.

The unparenthesized biomass and exchange values describe the constant-biomass reference calculation ($B_0 = B_{\max} = 1$, $\mu = 0$, and complete cell and protein retention at exchange). Parenthetical values specify the growth-aware protocol calculation: growth rate $\mu = 1.7\ \mathrm{h}^{-1}$, initial and stimulation biomass $B_{\mathrm{stim}} = 0.05$ OD, and retention $D_{\mathrm{wash}} = 1/4$ at later washes. In both cases extracellular trace is cleared at each exchange by medium replacement and carried through the reset to $B_{\mathrm{stim}}$ in the model. Thresholds therefore depend on the biomass present when the loop must hold its state.

The physical doses in the table are the applied concentrations. Their normalized counterparts

Table S1. Model inputs by source. The half-response scales $\kappa$ are fitted to the measured promoter responses (Fig. S1). The doses and phase durations are the applied protocol. The design point is the setting of the model trajectories. Parentheses give the growth-aware protocol values where they differ from the constant-biomass reference.

| Parameter | Value | Parameter | Value |
|---|---|---|---|
| *Fitted to measured responses* | | | |
| $\kappa_{\mathrm{las}}$ | $10^{-8}$ M | $\kappa_{\mathrm{lac}}$ | $7 \times 10^{-6}$ M |
| $\kappa_{\mathrm{lux}}$ | $5 \times 10^{-10}$ M | $\kappa_{\mathrm{tet}}$ | $2.7 \times 10^{-8}$ M |
| *Applied protocol* | | | |
| $s$ | 10; $C_{12}$ dose = 100 nM | $T_{\mathrm{active}}$ | 6 h |
| $i$ | 100/7; IPTG dose = 100 µmol L$^{-1}$ | $T_{\mathrm{wash}}$ | 2 h |
| $a$ | 1000/27; aTc dose = 1 µmol L$^{-1}$ | | |
| *Design point of the model trajectories* | | | |
| $\omega$ | 2, write capacity | $\delta$ | 2 h$^{-1}$ |
| $\xi$ | 9, feedback gain | $\mu$ | 0 (1.7) h$^{-1}$ |
| $\eta_\infty$ | 6, saturated erase load | $B_0$ | 1 (0.05) OD |
| $\rho$ | 1, trace/protein turnover ratio | $B_{\max}$ | 1 OD |
| $D_{\mathrm{wash}}$ | 1 (1/4), retained fraction | $B_{\mathrm{stim}}$ | not used (0.05 OD) |

are obtained by dividing each concentration by the corresponding module half-response scale:

$$s = \frac{[\mathrm{C}_{12}]}{\kappa_{\mathrm{las}}}, \qquad i = \frac{[\mathrm{IPTG}]}{\kappa_{\mathrm{lac}}}, \qquad a = \frac{[\mathrm{aTc}]}{\kappa_{\mathrm{tet}}}, \qquad x = \frac{[\mathrm{C}_6]}{\kappa_{\mathrm{lux}}}. \tag{S1}$$

The dose ratios are $s = 10$, $i = 100/7$, and $a = 1000/27$; each is above its half-response concentration. A value of one on a normalized axis is one half-response concentration.

The four dimensionless groups of Sec. III are

$$\omega = \frac{\beta\alpha_A}{\delta\delta_x}, \qquad \xi = \frac{\beta\alpha_M}{\delta\delta_x}, \qquad \eta_\infty = \frac{\gamma\alpha_E}{\delta\delta_x}, \qquad \rho = \frac{\delta_x}{\delta}, \tag{S2}$$

and the design point is $\omega = 2$, $\xi = 9$, $\eta_\infty = 6$, $\rho = 1$. Each group compares a production or removal process with the turnover of intracellular synthase and extracellular trace. The individual rates set how quickly states change, while the groups determine whether writing, feedback, and erasure are strong enough relative to loss. The groups are dimensionless under the convention that biomass represents culture volume and molecular variables are culture totals.

The write capacity $\omega$ is the maximum AND-module contribution to trace production when both conditioning inputs saturate. It describes the drive available during paired stimulation and is not itself a switching threshold.

The feedback gain $\xi$ measures how strongly the MEMORY population can replenish trace as a function of existing trace. In the reduced no-write, no-eraser system, bistability first appears at gain two, so $\xi = 9$ leaves a wide separation between the dark and bright steady states. During a wash, however, lower retained biomass reduces effective production and can remove that separation; this is why biomass and exchange parameters matter alongside $\xi$.

The saturated erasure load $\eta_\infty$ is the maximum AiiA-mediated trace removal relative to spontaneous trace loss, reached when the eraser input is saturating. Its time-dependent value follows the AiiA pool:

$$\eta = \frac{\gamma A}{\delta_x}, \qquad \frac{\mathrm{d}\eta}{\mathrm{d}\tau} = \eta_\infty e - \eta, \qquad e = \theta(a). \tag{S3}$$

Thus $\eta_\infty$ is the eventual load under full induction, not the instantaneous load at the start of erasure; finite enzyme accumulation makes duration relevant. Finally, $\rho = 1$ means that extracellular trace and intracellular synthase decay at the same rate.

### S2. PROMOTER-RESPONSE CALIBRATION AND FITTING

The six panels of Fig. S1 summarize experimental calibration of the promoters used in the consortium. Panels (a)–(d) give the single-input responses and panels (e) and (f) the two-input responses of the AND and RECALL gates, each measured in three independent cultures per input condition. For a direct GFP reporter, the promoter-negative background is subtracted and the remainder scaled by the dynamic range between that background and the saturating response. For culture $r$ at concentration $c_j$, the normalized measurement is

$$y_{rj} = \frac{F_{rj} - F_{\mathrm{auto}}}{F_{\mathrm{sat}} - F_{\mathrm{auto}}}. \tag{S4}$$

The dose-response model of Eq. (S5) estimates the basal fraction $\ell$ and the half-response concentration $\kappa$ at the Hill coefficient $n = 2$ that Eq. (2) assumes throughout the model:

$$f(c; \ell, \kappa) = \ell + (1 - \ell)\frac{c^2}{\kappa^2 + c^2}. \tag{S5}$$

Here $f(0) = \ell$, $f(\kappa) = (1 + \ell)/2$, and the high-dose limit is one. Fixing $n$ keeps the fitted scales in the units the model is written in; releasing it is a check on that assumption rather than part of the calibration, since a fitted $n \neq 2$ would change the fold conditions of Sec. III C and not merely the scales.

The response is fitted jointly across concentrations and independent cultures by nonlinear least squares,

$$(\hat{\ell}, \hat{\kappa}) = \underset{0 \le \ell < 1,\, \kappa > 0}{\operatorname{argmin}} \sum_j \sum_r w_{rj} \left[ y_{rj} - f(c_j; \ell, \kappa) \right]^2. \tag{S6}$$

The response is fitted to the three culture-level measurements at each dose. Weights are $w_{rj} = 1/\sigma_j^2$, with $\sigma_j$ the between-culture standard deviation at dose $c_j$.

The single-input response curves are fitted to the four measured module responses; the gated response data are compared with the product prediction from those fits. The half-response scales define the normalized inputs of the model, and the fitted basal fractions are annotated in Fig. S1.

For AND and RECALL both promoter inputs are calibrated and the separable-gate assumption tested. If $c$ is the HSL input and $q$ is held IPTG, the predicted gated response is

$$f_{\mathrm{gate}}(c, q) = f_{\mathrm{HSL}}(c; \ell_H, \kappa_H) f_{\mathrm{IPTG}}(q; \ell_I, \kappa_I). \tag{S7}$$

The three held-IPTG series in panels (e) and (f) test the predicted change in gate contrast. Deviations from the product surface indicate promoter interaction or other nonseparability and should be fitted with a joint response surface rather than forced into Eq. (S7). The pLux and pLas–*lacO* modules produce LuxI and hence extracellular lactone rather than fluorescence, so they are read through the TOP10 pLux–*gfp* sensor. Until that sensor is calibrated against lactone standards in matched medium, their scales are relative to the sensor and not to a concentration.

The fitted half-response scales enter Table S1 and define the normalized inputs of every rule in Secs. III C and III D. The measured fits and their three-culture replication are the empirical calibration used here.

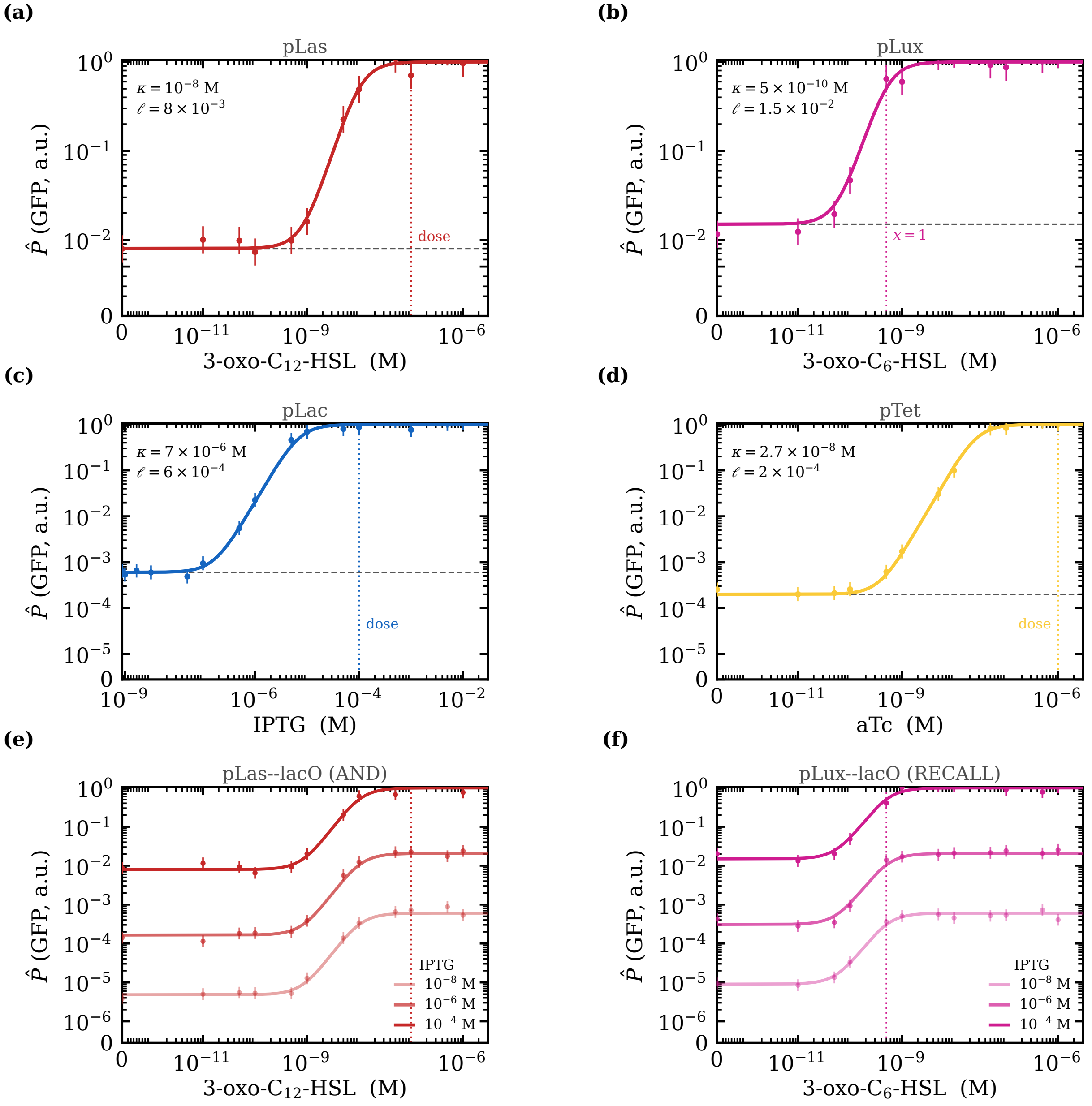


Figure S1. Experimental promoter-response calibration. Panels (a)–(d) show measured single-input pLas, pLux, pLac, and pTet response profiles from three independent cultures per ligand concentration, normalized to each module's own maximum; fitted half-response scales $\kappa$ and basal fractions $\ell_j$ are annotated. Panels (e) and (f) show measured pLas–*lacO* AND and pLux–*lacO* RECALL gate responses from three independent cultures at each HSL concentration and held IPTG level. Solid curves show the product predictions from the single-input fits. Vertical dotted lines mark the protocol doses; in panel (b), the line marks the normalized signal $x = 1$. Concentrations are in M.
